# Root economics spectrum and construction costs in Mediterranean woody plants: the role of symbiotic associations and the environment

**DOI:** 10.1101/2020.06.08.139691

**Authors:** Enrique G. de la Riva, Iván Prieto, Teodoro Marañón, Ignacio M. Pérez Ramos, Manuel Olmo, Rafael Villar

## Abstract

1. Quantifying the functional variation of fine root traits and their interactions with symbiotic organisms is an uprising research topic to understand the overarching trade-off between maximizing resource acquisition or conservation (root economics spectrum -RES-). The currency of root traits economy is based on the carbon required to construct them; however, we lack a clear understanding of this question.
2. Our main aim was to quantify construction costs (CC) of fine roots (<2 mm) and their underlying components (concentration of carbon, minerals and organic nitrogen) in 60 Mediterranean woody species with contrasting symbiotic association types (ectomycorrhizas, arbuscular and ericoid mycorrhizas and N-Fixing nodules). We examined (1) if the CC depended on the symbiotic type, and if they were associated with morphological traits; (2) the relation of each component of the CC with the root structure for each symbiotic type; and (3) whether soil water and nutrient availability determined differences in CC across sites and symbiotic types.
3. The multivariate analysis of root traits showed a main plane of covariation accordingly to the RES expectations, with contrasting symbiotic types at both ends of the spectrum. We found a significant relationship between root CC and RES; interestingly the slopes of this relationship differed among symbiotic types, which was mainly due to the different role that each specific components of CC plays across them. In addition, independently of the symbiotic type, the CC decreased linearly with the nutrient availability and quadratic with the water availability.
4. *Synthesis*. Our study demonstrates that woody plants have different strategies in their root carbon investment, which depend on their position along the economics spectrum (RES) and on their main symbiotic association preference. The intrinsic components of the cost of root production varied across species with contrasting symbiotic associations, pointing to a trade-off between structural and metabolic compounds. We also found that root CC are strongly modulated by soil resource availability, following a non-linear pattern with water availability. Thus, CC shifts from high to low at the most arid sites, which points to a strong role of symbiotic associations in this shift.

## INTRODUCTION

The functional variation of fine root traits and their interactions with symbiont organisms is an uprising research topic to better understand the large diversity of plant belowground uptake strategies across the world (Kong et al. 2019; Bergmann et al. 2020; Shi, Li, Zhu & Wang 2020). Leaf traits covary across plant species following an overarching trade-off between maximizing resource acquisition and productivity or maximizing resource conservation at a productivity cost, the so-called leaf economics spectrum (LES, Wright et al. 2004). Seeking similar patterns related with a resource uptake trade-off on fine roots (< 2 mm diameter), several studies have suggested that the economics spectrum theory can also be applied to root uptake strategies (root economics spectrum -RES-; Prieto et al. 2015; Roumet et al. 2016; de la Riva et al. 2018a; Bergmann et al. 2020; Marañón et al. 2020). However, despite the numerous studies made by plant ecologists, especially in the last years, the covariation of root traits according with resource economics is still under debate (Chen, Zeng, Eissenstat, & Guo 2013; Kramer-Walter et al. 2016; Weemstra et al. 2016; Kong et al. 2019; Delpiano, Prieto, Loayza, Carvajal, & Squeo 2020). Alternatively, many authors suggest the existence of a multidimensional space of root trait variation where symbiotic associations may be playing a key role (Navarro-Fernández et al. 2016; Erkan, McCormack, & Roumet 2018; Bergmann et al. 2020).

The LES is based on the main idea that tissue construction is constrained by a fundamental trade-off between fast growth rate (or resource capture) and high survival (or resource conservation), which is well-represented along environmental resource gradients (Wright et al. 2004; Reich 2014). Assuming a functional similarity between leaves and roots, it would be expected that plant species growing in more productive environments have thinner roots with low tissue density as well as higher specific root length (SRL) or specific root area (SRA) with shorter lifespan, favouring strategies with a faster return of investments. By contrast, in environments with lower resource availability, opposite traits would be expected, resulting in roots with longer lifespan and slower return of investment (de la Riva et al. 2018a; Roumet et al. 2016). Mostly, the currency of root economics is the carbon required to construct fine roots that explore the soil for resource acquisition (Bergmann et al. 2020); for example, it is expected that SRA (i.e. the root area per unit biomass) reflects the available area for root uptake at a given biomass cost. The construction costs (CC) of roots is defined as the total amount of metabolic compounds required to construct all the chemical constituents of 1 g of root biomass (Villar, Robleto, De Jong, & Poorter 2006). This cost includes both the compounds consumed in respiration to supply reductant and ATP for energy-requiring processes in the biosynthesis of the tissue constituents, as well as those resources (such as carbon and organic and inorganic compounds) required to build the root (Penning de Vries, Brunsting & van Laar 1974; Poorter & Villar 1997). Nonetheless, inferences of morphological root traits as proxies of resource economics strategies based on CC remain rather speculative, because most of the suppositions regarding a relationship between root traits and resource economics have been based on trait correlations and/or their variation along resource gradients, while the actual cost and energy involved in developing the structure and function of these fine roots has been generally overlooked. That is, although root CC is frequently mentioned in the literature when as a factor explaining the variation in root trait values along the resource gradient by the economics spectrum theory, few studies have actually addressed this question experimentally using comparative analyses (Martínez, Lazo, Fernández□Galiano, & Merino 2002; Villar et al. 2006; Roumet et al. 2016).

One key strategy to enhance plant nutrient acquisition is the association of roots with symbiotic microorganisms that increase nutrient uptake (Wang, Pan, Chen, Yan & Liao 2011). These symbiotic associations imply root morphological changes (i.e. changes in root trait values) that could potentially decouple the relationships among root traits commonly defined by resource economics (e.g. Weemstra et al. 2016; McCormack et al. 2017; Bergmann et al. 2020). In this regard, Bergmann et al. (2020) have recently suggested an alternative gradient of root specialization, which ranges from roots with short leaf lifespan and traits associated to a fast resource acquisition for an efficient soil exploration (e.g. high SRL and low tissue densities), the ‘do-it-yourself’ strategy, to roots with a higher inversion of carbon to outsource resource acquisition via symbiotic associations to enhance nutrient acquisition. As the authors highlight, such ‘outsourcing’ strategy has morphological consequences for the roots; for example, some plant species develop thick roots (i.e. high diameter) with a thick cortex but a low root tissue density that increases the affinity for mycorrhizal fungi (Kong et al. 2014). Other species from the Fabaceae family, known to have leaf traits associated to fast growing strategies (Powers & Tiffin 2010), develop complex cluster roots with rhizobial N_2_ fixing bacteria, enhancing the resource uptake needed to compensate for the fast rates of respiration required to rapid root growth (Shane, Cawthray, Cramer, Kuo, & Lambers 2006; Li, Zeng, & Liao 2016). These types of symbioses may allow plants with thick, light roots to have an acquisitive strategy and grow in resource-limiting environments because symbionts increase the soil exploration capacity and enable a faster acquisition of nutrients and water (Kramer-Walter & Laughlin 2017). These types of strategies allow roots to increase their resource uptake with low values of tissue density, enhancing a faster pay back of the construction cost when environmental conditions become favourable; e.g. in Mediterranean environments the development of proteoid roots has been observed to fluctuate seasonally (Shane et al. 2006). However, there is not unanimous agreement in the mechanisms underlying the cost of these root modifications, and thus, little progress has been made in understanding the underlying role of the symbiotic associations over the root morphological remodeling in relation to resource use strategies (McCormack et al. 2017).

The main aim of the present study is to quantify root construction costs and their underlying components (concentration of carbon, minerals and organic nitrogen) in 60 Mediterranean woody species with contrasting symbiotic association types. These species are a subset from a previous study with 80 woody Mediterranean species where we observed that root morphological traits covaried as expected by the RES theory (de la Riva et al. 2018a). Based on current knowledge, we might expect that the roots of the acquisitive species differ from the conservative ones in the structural and chemical characteristics, because low resource availability would lead to greater investment in costly structural compounds (Martínez et al. 2002; Villar et al. 2006). Thus, the energetic costs of tissue construction of roots might be expected to be higher in species from unproductive habitats than in those inhabiting more favourable environments, showing strong covariation between the RES and the CC. However, in such approach, the main difficulties for interpretation arise from the potentially large impacts of mycorrhizal fungi and rhizobial bacteria on root traits, which may confound the covariation among the morphological traits and the energetic costs of tissue construction (Li et al. 2016; Weemstra et al. 2016). To clarify these possible sources of variation, we included into our analyses four symbiotic association types (ectomycorrhizas, arbuscular mycorrhizas, ericoid mycorrhizas and N-Fixing nodules) in order to assess the impact of the symbiotic types over the observed RES patterns in Mediterranean woody species, and their influence on the root CC. Specifically, we firstly assessed if the root CC of fine roots was associated with morphological traits, by quantifying whether the covariation of root traits along the RES was related to the intrinsic cost of root tissues; in addition we tested if that relation depended on the symbiotic type. Secondly, we assessed the relationship of each component of the construction cost (concentration of carbon, minerals and organic nitrogen) with the RES, for each symbiotic type, to elucidate the role of each particular component on root morphological variation. Thirdly, we quantified whether soil water and nutrient availability also determined differences in construction costs across sites.

## MATERIAL AND METHODS

### Study area and species selection

We selected the most abundant and representative woody species from Mediterranean forests and shrublands across seven sites in southern Spain. These sites spanned a wide gradient of soil water and nutrient availability, ranging from riparian forest in mountain systems (Las Tonadas, Las Tonadas canyon and Las Navas, in Dehesas de Sierra Morena Biosphere Reserve) to coastal shrublands (Monte Negro and Monte Blanco, Doñana National Park), sub-humid forests (Los Alcornocales Natural Park) and arid shrublands (Cabo de Gata-Níjar Natural Park, Appendix S1, Supporting Information). At each study site, we selected four to six individuals of each woody species, 60 species in total, and measured their fine (<2 mm) root functional traits. Most species were only present in one site, whereas others were common to more than one site making a total of 73 site × species combinations (Appendix S2).

Plant species were classified into four categories according to their main symbiotic association type: ectomycorrhizal species (EcM), defined by the presence of a Hartig net and mantle (Brundrett & Tedersoo 2018); arbuscular mycorrhizal species (AM), defined by the presence of an intracellular surface mycelia colonization with arbuscules and vesicles; ericoid mycorrhizal species (ErM), defined by the presence of an intracellular surface mycelia colonization with interwoven coils (Brundrett & Tedersoo 2018); and N-fixing species (N-Fix), which are species from the Fabaceae family which develop dense clusters of lateral roots and/or nodes containing rhizobia (i.e. N-fixing bacteria, Li et al. 2016). We assigned a mycorrhizal status to each species based on our own measurements for the species present at the Sierra Morena sites (Navarro-Fernández et al. 2016), and from the FungalRoot database for the species located in the remaining sites (Soudzilovskaia et al. 2020) (see Appendix S2). Eight species could not be assigned using these two sources, so in order to complete the dataset we filled the gaps following Bergmann’s approach (Bergmann et al. 2020): 1) almost all plant species have one type of dominant mycorrhizal association and 2) the mycorrhizal association type is usually very conserved within a monophyletic genus and often within a family. Therefore, these species were assigned with the mycorrhizal association type of the closest phylogenetically sister species.

### Root trait measurements

Fine roots (< 2 mm in diameter) were sampled in four individuals per species and site by excavating the top 20–30 cm of soil close to the plant basal stem. Excavated roots were washed using a 0.5 mm sieve, weighed to determine their fresh mass (FM, g) and scanned at 1200 dpi. Roots were then dried at 60□C during at least 48 h, and weighed to obtain their dry mass (DM, g). Root length (L, cm), diameter (Rdi, mm), area (A, cm^2^) and volume (V, cm^3^) were obtained by analyzing the scanned root samples with specific software (WinRHIZO 2009, Regent Instruments Inc., Quebec, Canada). Specific root length (SRL, m g^−1^) and specific root area (SRA, m^2^ Kg^−1^) were determined as (L/100)/DM and (A/10000)/DM, respectively. Root dry matter content (RDMC, g g^−1^) and root tissue mass density (RTD, g cm^−3^) were determined as DM/FM and DM/V, respectively. Five root traits -SRL, SRA, RDMC, RTD and Rdi-were selected for statistical analysis (see below).

All root trait measurements were carried out according to the criteria and methodology defined by Pérez□Harguindeguy et al. (2013). For a more detailed protocol of root sample harvesting and trait measurements see de la Riva et al. (2016a).

### Construction cost measurements

The construction cost of each species was determined in a mixture of roots from four individuals. For each species we measured the concentration of C, minerals and organic N, following the method of Vertregt & Penning de Vries (1987) as modified by Poorter (1994). An exact description of the procedures followed for the chemical analyses and the subsequent calculations is given in Poorter & Villar (1997).

### Soil physical and chemical properties

The soil was characterized in each of the seven sampling sites. We collected four soil samples in each site from the top 20 cm using a soil auger; within that soil depth is where nutrient uptake mostly occurs (Jobbágy & Jackson 2001). In the laboratory, soil samples were air-dried, crushed and sieved at 2 mm, and soil organic matter, nutrients (P, N, Mg, K and Ca) and potential water availability (PWA) were determined using standard soil methods (Sparks 1996). Soil organic matter was determined by the Walkley and Black method, and total N was determined by Kjeldahl digestion. Available P was estimated by the Olsen method, while available Ca, K and Mg were extracted with 1 M ammonium acetate and determined by atomic absorption spectrophotometry. To characterize the soil water availability at each site, we used the potential soil water availability of each site (PWA, see de la Riva et al. 2018a for more details).

### Data analyses

At a preliminary step, to obtain an overview of the multidimensional spectrum of root trait variation, a principal component analysis (PCA) was performed with the five root traits for the 73 observations (mean values per species x site). We assume that the main variation trend (PC1) represents the root economics spectrum (RES, de la Riva et al. 2018a). Following, to assess the segregation of plant species according to their symbiotic association type (EcM, AM, ErM and N-Fix) along the main variation trends of traits, we used a one-way ANOVA with the PCA scores of the first and second components as the dependent variables and the symbiotic association type as the factor. Post-hoc Tukey tests were performed to check the significance of the pairwise differences between symbiotic association types.

Regarding the root construction costs (CC), we firstly tested differences in CC among symbiotic association types with a one-way ANOVA. Next, general linear models (GLM) were used to test the relationships between root construction costs and the RES (i.e. PC1 scores), as well as between root CC and each of the morphological root traits considered in this study. We included the symbiotic association type in the models to test differences between the slopes of the relationships across symbiotic types using an analysis of covariance ANCOVAs (Kurokawa & Nakashizuka 2008). Once observed that there were large differences among symbiotic association types in the slopes of relationships between CC and the RES, we tested the influence of each component of the construction cost independently for each symbiotic type [C: concentration of C, Min: minerals and OrgN: organic N) using multiple regression analyses (except within ErM for which we only had four observations). For this purpose, we conducted a decomposition analysis based on the Sum of Squares (SS) of each component. The SS can be decomposed into the amount of variability explained by individual terms of the model (SS_C_, SS_Min_ and SS_OrgN_) and by the unexplained variability (SS_Error_) given SS_RES_ = SS_C_ + SS_Min_ + SS_OrgN_ + SS_Error_. For example, the percentage of explanation of the root carbon concentration (C) on the variation of root construction cost would be given by (SS_C_/SS_RES_) × 100. To obtain the variability explained by each component without covariation, we used the type III sum of squares.

In addition, we used general lineal models (GLM) to assess the relationship between soil physical and chemical properties, and root construction costs. Firstly, to reduce the number of variables characterizing the resource environmental gradient and their collinearity, a PCA was performed with the abiotic variables measured in this study: soil nitrogen (N) and organic matter (OM), availability of nutrients (P, Mg, K and Ca), and potential water availability (PWA) (Appendix S3). Secondly, we conducted a linear regression with quadratic components (Y_i_ = β0 + β_1_X_i_ + β_2_X_i_^2^ + ε_i_), with the CC as dependent variable and the scores of the first axis of the abiotic PCA as independent factor, which represents a gradient of nutrient and water availability.

In order to control for potential phylogenetic effects on construction costs and trait covariation, all the relationships between morphological traits and construction costs were compared with the results obtained by fitting a phylogenetic generalized least squares model (pgls). To this end, we used the pgls function of the caper package (Orme 2013), which addresses phylogenetic non-independence among species by incorporating covariance between taxa into the calculation of the estimated coefficients. The phylogenetic tree (Appendix S4) was obtained with the comprehensive Angiosperm species-level phylogeny from Zanne et al. (2014), as updated by Qian & Yin (2016), which is included into the R package ‘S. PhyloMaker’ (Qian & Yin 2016). The distance of the few species that were not found in the PhytoPhylo database were supplanted by the distance of the closest species of the same genus found in the mega-phylogeny tree (de la Riva et al. 2019).

All calculations and statistical analyses were performed with the R software (v 2.15.3, R Core Team 2019).

## RESULTS

### Root construction costs and root economics spectrum as a function of plant symbiotic type

The principal component analysis showed that covariation among fine root traits was represented by two independent dimensions, encompassing a bi-dimensional plane with a cumulative explanatory power of 94% of all root trait variation (Fig. 1). The first principal component (PC1) represented the variation along the root economics spectrum (RES) with 62.9% of the total variation explained (Fig. 1). This first axis (PC1) scores were negatively associated to SRA and SRL and positively to RDMC and root tissue density. The second principal component explained 31.2% of the overall variance and was represented mainly by the trade-off between root tissue density (negative scores) and root diameter (positive scores). Interestingly, plant species with contrasting symbiotic association types occupied separate spaces within the first PCA axis, potentially representing different resource-use strategies (F_3,69_=4.84, *P*=0.004); thus, fine roots of AM species had scores associated to acquisitive strategies (i.e. negative scores) in opposition to fine roots from EcM species that had scores associated to more conservative strategies (Fig. 1). Fine roots from Nitrogen fixing (N-Fix) and ErM species had intermediate scores and were distributed along all this first axis.

**Figure 1.**
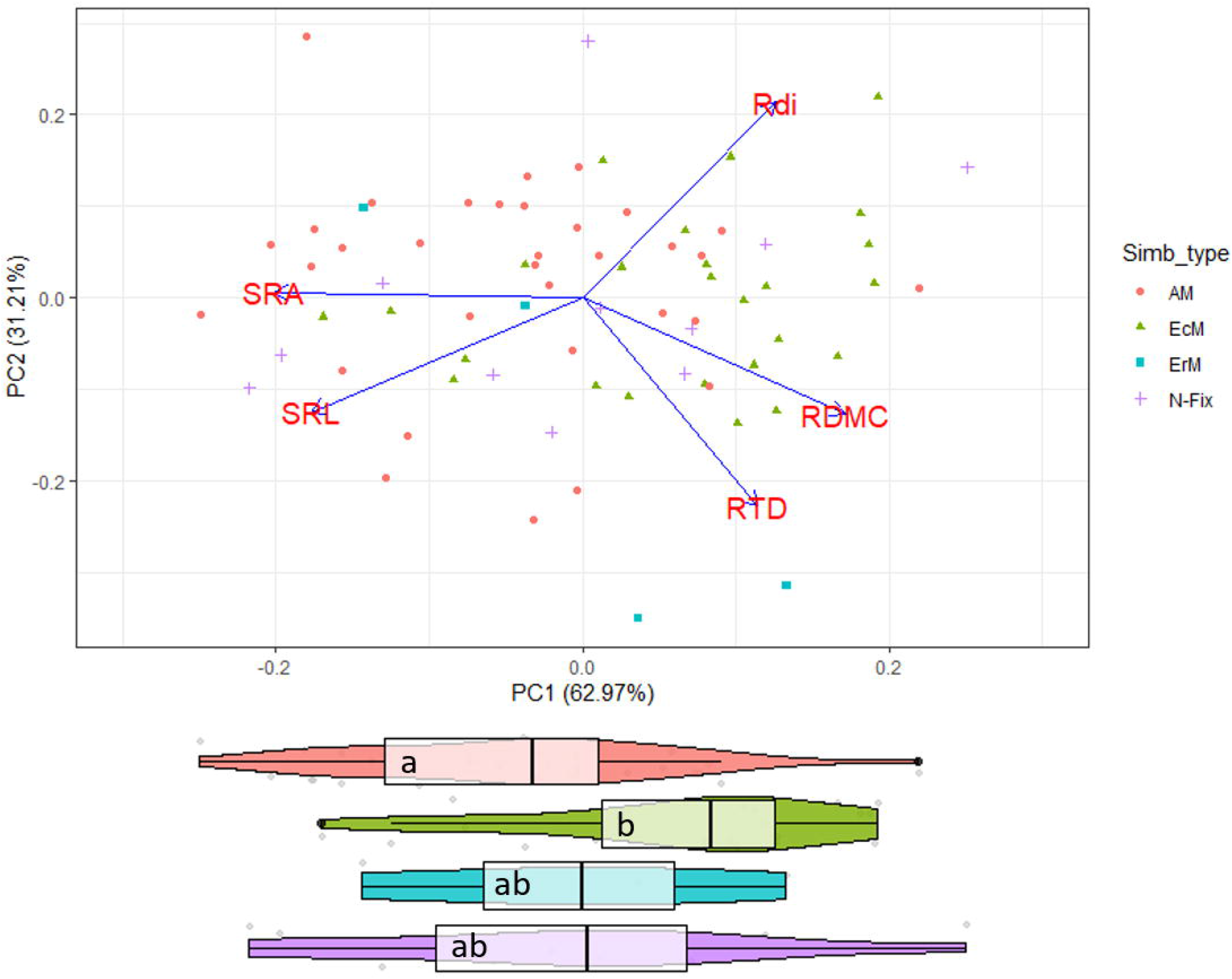
Plot of the first and second axes of the principal component analysis (PCA) performed with five root morphological traits (SRL: specific root length; SRA: specific root area; RDMC: root dry matter content, RTD: root tissue density and Rdi: root diameter) and 60 Mediterranean woody species growing along a gradient in resource availability. Each dot represents a mean value for each individual species and site (73 species × site combinations), and different colours represent different symbiotic association types: Arbuscular mycorrhizal species (AM) in red; Ectomycorrhizal species (EcM) in green color; Ericoid mycorrhizal species (ErM) in blue, and N_2_ fixing species (N-Fix) in purple. Violin plots with median values and 1^st^ and 3^rd^ quartiles (outside box lines) are included in the bottom; different letters indicate significant differences in mean PC1 score values among species with contrasting symbiotic association (Tukey post-hoc test, *P* < 0.05).

Although we did not find any significant differences in root construction cost between symbiotic association types (F_3,69_=0.44, *P*=0.724), we found a significant positive relationship between root construction cost (CC) values and their scores along the first PCA axis describing the root economics spectrum (RES) (F_3,65_=6.50, *P*<0.001; Fig. 2), indicating that conservative strategies imply greater root construction costs than acquisitive strategies. Interestingly, the slope of the relationship differed for each symbiotic association type (ANOVA slope test, F_3,65_=6.50, *P*<0.006; Fig.2). In addition, we found a positive significant relationship between root construction costs and the RES for AM (F_1,31_=7.67, *P*<0.009), N-fix species (F_1,9_=16.79, *P*<0.002), and marginally significant for ErM (F_1,2_=7.45, <0.1), but no significant relationship for EcM species. The second PCA axis was also negatively related with fine root construction costs (F_1,71_=7.78, *P*=0.006) but no significant interactions within symbiotic status were observed (Appendix S5).

**Figure 2.**
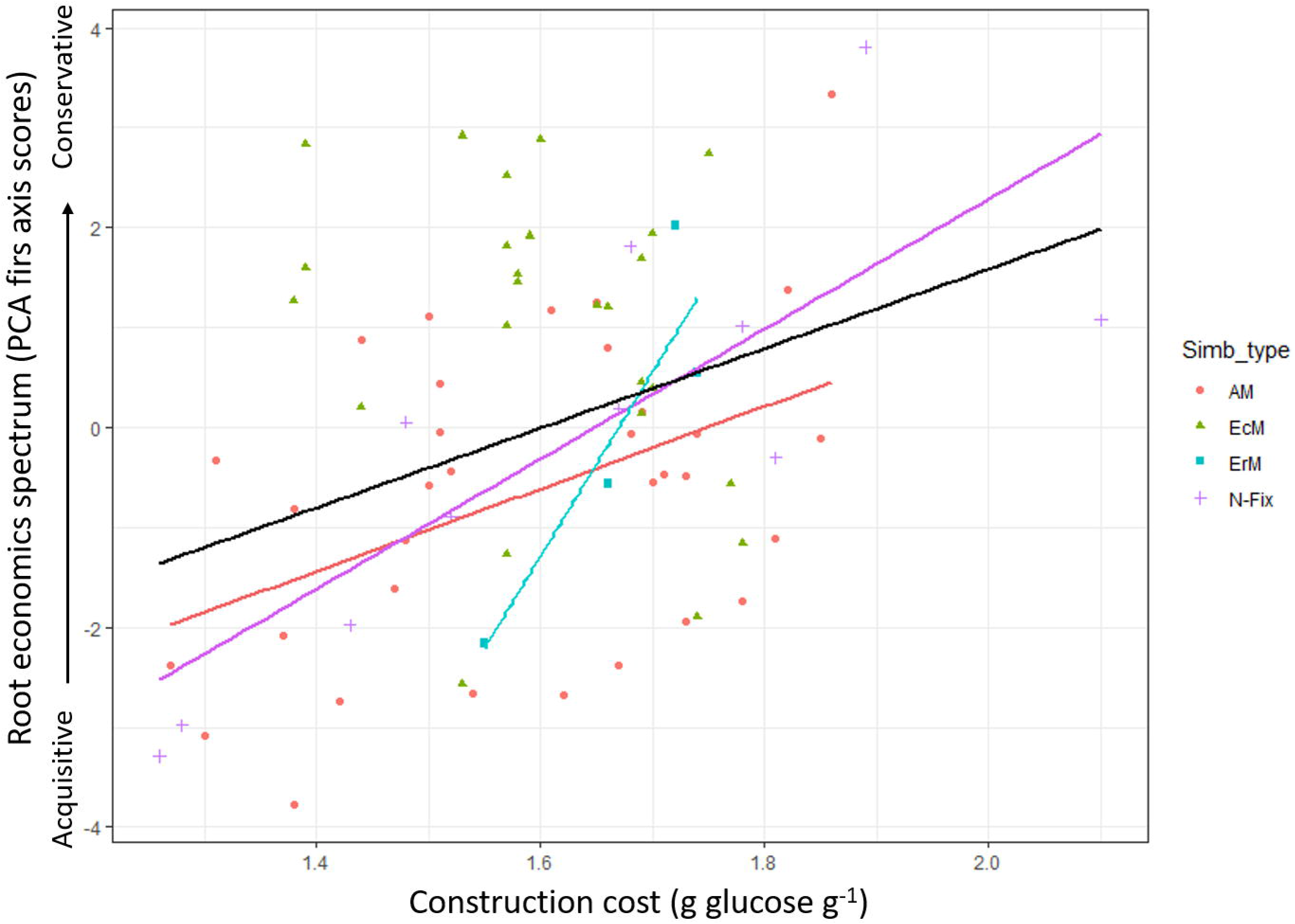
Relationships between root construction costs and the first PCA axis (PC1) depicting the root economics spectrum (RES). The global relashionship (considering all data, black line) and for each type of simbiotic association type (when significant) are shown. Abbreviations: Arbuscular mycorrhizal species (AM); Ecto mycorrhizal species (EcM); Ericoid mycorrhizal species (ErM) and N-fixing species (N-Fix).

When analysing the relationships between root construction cost and each morphological root trait separately we found that construction cost was negatively related with SRA (and marginally with SRL), while positively related with RTD and RDMC (Table 1). Most of these relationships with single traits showed significant interactions between construction costs and symbiotic association type. Results of the relationships were consistent after accounting for phylogenetic effects (PGLM in Table 1), which suggests that these relationships are phylogenetically independent.

**Table 1.**
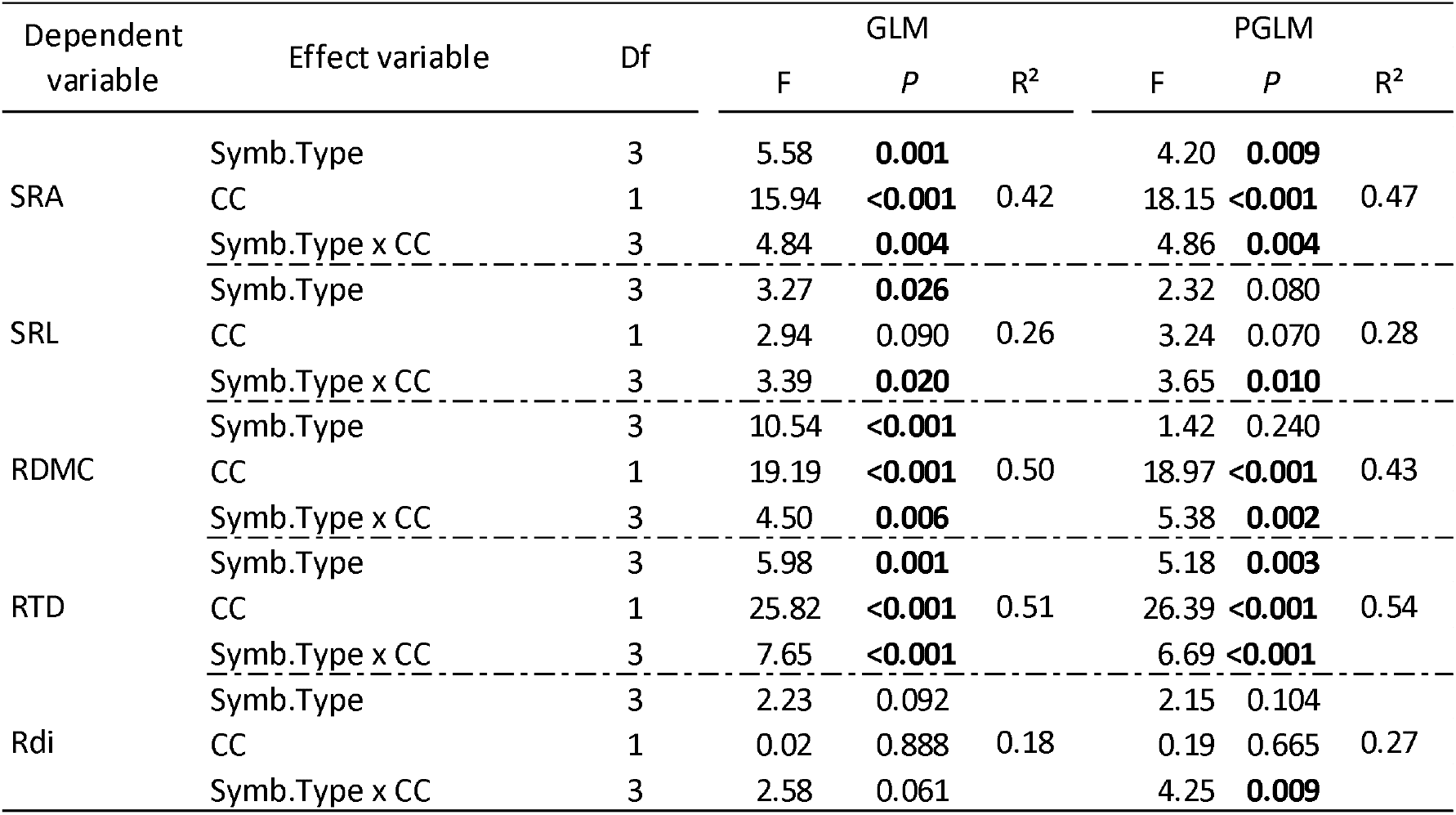
Results of the relationships (general linear models: GLM, and phylogenetic generalized linear models: PGLM) between morphological root traits (SRA: specific root area; SRL: specific root length; RDMC: root dry matter content, RTD: root tissue density and Rdi: root diameter) and root construction costs (CC), and differences among slopes for symbiotic association types (Arbuscular mycorrhizas, Ecto mycorrhizas, Ericoid mycorrhizas and N-fixing species) in 60 Mediterranean woody species growing along a gradient in resource availability. The degrees of freedom (Df), F-values (F), significance (*P*-value) and R^2^ of the models are shown.

| Dependent variable | Effect variable | Df | GLM |  |  | PGLM |  |  |
| --- | --- | --- | --- | --- | --- | --- | --- | --- |
|  |  |  | F | P | R<sup>2</sup> | F | P | R<sup>2</sup> |
| SRA | Symb.Type | 3 | 5.58 | **0.001** |  | 4.20 | **0.009** |  |
|  | CC | 1 | 15.94 | **<0.001** | 0.42 | 18.15 | **<0.001** | 0.47 |
|  | Symb.Type x CC | 3 | 4.84 | **0.004** |  | 4.86 | **0.004** |  |
| SRL | Symb.Type | 3 | 3.27 | **0.026** |  | 2.32 | 0.080 |  |
|  | CC | 1 | 2.94 | 0.090 | 0.26 | 3.24 | 0.070 | 0.28 |
|  | Symb.Type x CC | 3 | 3.39 | **0.020** |  | 3.65 | **0.010** |  |
| RDMC | Symb.Type | 3 | 10.54 | **<0.001** |  | 1.42 | 0.240 |  |
|  | CC | 1 | 19.19 | **<0.001** | 0.50 | 18.97 | **<0.001** | 0.43 |
|  | Symb.Type x CC | 3 | 4.50 | **0.006** |  | 5.38 | **0.002** |  |
| RTD | Symb.Type | 3 | 5.98 | **0.001** |  | 5.18 | **0.003** |  |
|  | CC | 1 | 25.82 | **<0.001** | 0.51 | 26.39 | **<0.001** | 0.54 |
|  | Symb.Type x CC | 3 | 7.65 | **<0.001** |  | 6.69 | **<0.001** |  |
| Rdi | Symb.Type | 3 | 2.23 | 0.092 |  | 2.15 | 0.104 |  |
|  | CC | 1 | 0.02 | 0.888 | 0.18 | 0.19 | 0.665 | 0.27 |
|  | Symb.Type x CC | 3 | 2.58 | 0.061 |  | 4.25 | **0.009** |  |

### Components of the root construction costs and symbiotic types

The partitioning analysis of the total variability among the different components of root construction costs within the RES (PC1) demonstrated that the role of each component varied across symbiotic association types (Fig. 3). For EcM and N-Fixing species, the relationship between the RES and construction costs was explained in a large proportion by the variation in the concentration of root carbon (20% and 23%, respectively), mineral concentration (29%, for EcM species), while organic nitrogen was the most important component for N-fix species (22%). In contrast, for AMF species, the relationship between construction cost and the RES was poorly explained by the root C variation among species (1%; Fig. 3), being mostly explained by variations in organic nitrogen (14%) and the concentration of minerals (9%).

**Figure 3.**
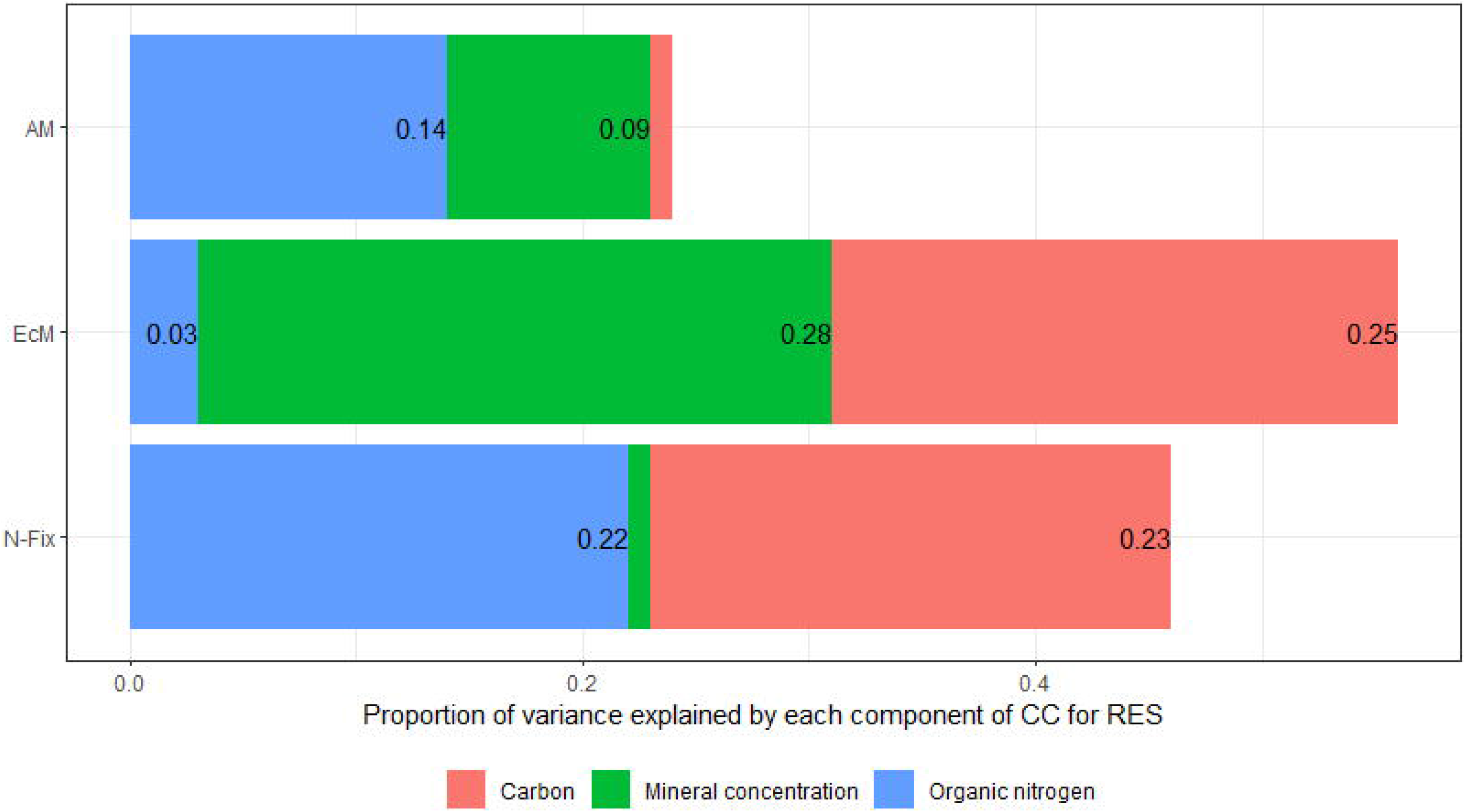
Decomposition of the total variability explained by each component of fine root construction costs (concentration of minerals: Conc_Min; organic nitrogen: N_org; and carbon: C) on the first axis of variation of PCA depicting the RES, for each symbiotic association type: Ecto mycorrhizal species (EcM), Arbuscular mycorrhizal species (AM) and N-fixing species (N-Fix). ErM species were not included in this analysis due to their reduced number of samples (n=4).

**Figure 4.**
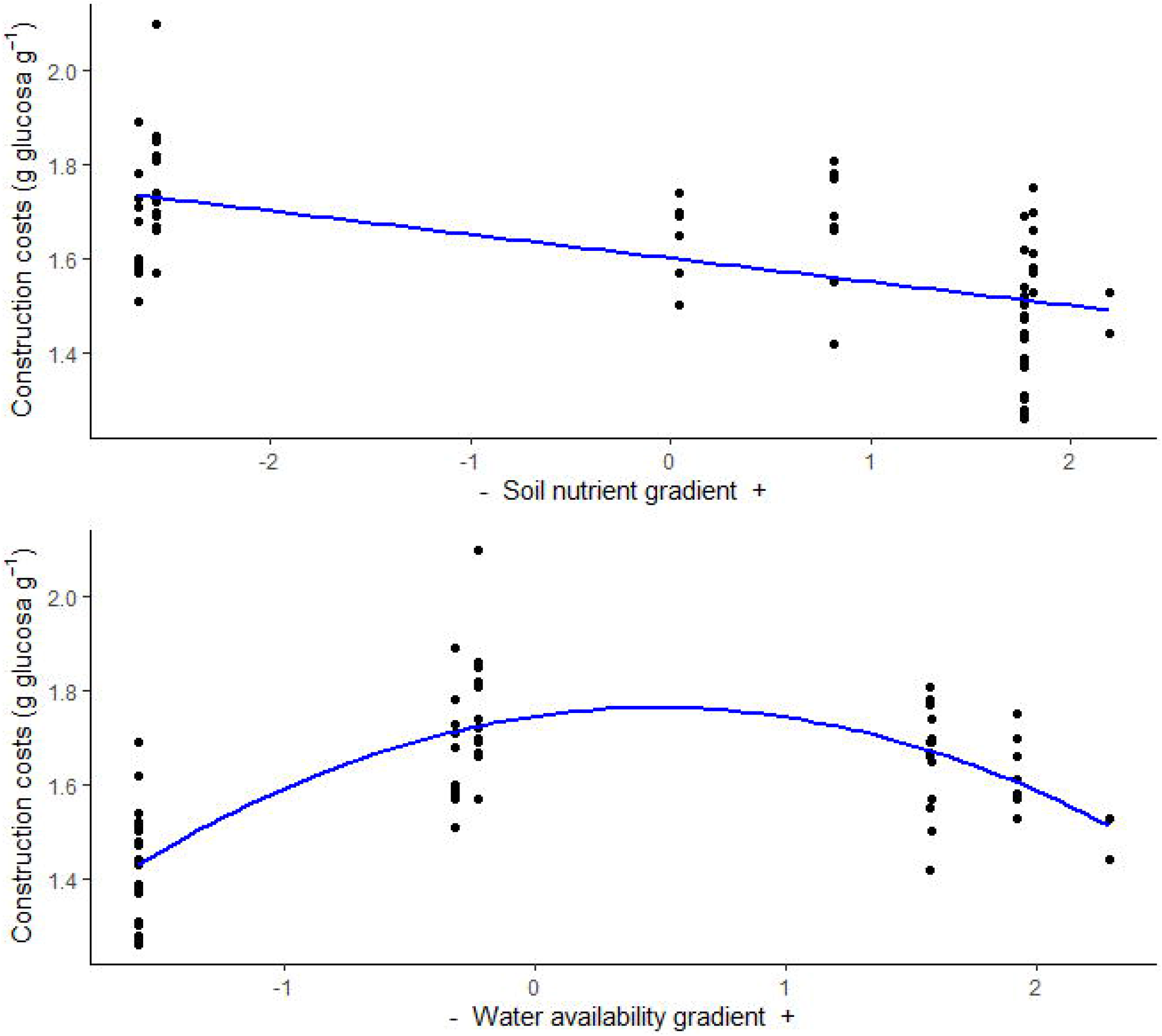
Relationships between root construction costs and the two main PCA axes representing a gradient in soil nutrient availability (PC1, R^2^= 0.33, *P* <0.001) and a gradient in soil water availability (PC2, R^2^= 0.56, *P* <0.001; quadratic relationships: Yi = β0 + β1Xi + β2Xi ^2^ + εi).

### Root construction costs along the soil resource gradient

The first two axes of the principal component analysis (PCA) for environmental properties explained 85.2% of the total variance (Appendix S3). The first PCA axis explained 55.7% of the total variance and represented a gradient of soil nutrient availability (concentration of soil nitrogen, magnesium, phosphorous, calcium and potassium and organic matter) while the second PCA axis (26.9 % of the variance) was mainly associated to differences in the potential water availability. A significant linear negative relationship was observed between the first PCA axis (nutrient availability) and fine root construction costs (*P*<0.001, R^2^=0.36); fine root construction costs were higher in species growing in nutrient poorer areas (i.e. negative scores of the resource gradient PCA, e.g. Doñana National Park) and decreased as we moved to most productive sites (e.g. Alcornocales Natural Park, positive scores of the resource gradient PCA). Contrastingly, the relationship between the second PCA axis and fine root construction costs was quadratic (*P*<0.001, R^2^=0.56); species from the Cabo de Gata Natural Park and the riparian areas, i.e. the most arid and most humid environments, respectively (Appendix S1), showed very low root construction cost values.

## Discussion

### Relationships between root construction costs and economics spectrum as a function of plant symbiotic type

The knowledge of the energy content of the constituents involved in establishing the structure and function of the root system is essential for the analysis of plant performance and ecosystem processes (Martínez et al. 2002; Laureano et al. 2013). In this study, we identified a set of covarying morphological root traits (i.e. SRA, RDMC and RTD) that revealed strong links with the total amount of metabolic compounds required to construct root biomass. In agreement with the root economics spectrum (RES) expectations, we demonstrated that lower construction costs were associated with higher foraging ability (i.e. higher values of SRA and SRL), a syndrome commonly associated with a fast return of investments. By contrast, higher construction costs were observed in the species with more conservative strategies (i.e. higher RDMC and RTD), which confers greater abundance of metabolic defense and repair complexes in the root tissues at the expense of higher metabolic machinery and higher maintenance costs (Laureano et al. 2008). Moreover, this relationship held within groups of species with different symbiotic associations, except for EcM. To our knowledge, this is one of the first studies reporting a relationship between the RES and root construction costs, expanding our knowledge on root functioning in woody species (shrubs, arborescent-shrubs and trees). Our results are in line with a previous study reporting a positive relationship between acquisitive strategies and higher root respiration rates in 62 herbaceous and 12 dwarf shrub species (Roumet et al. 2016). Despite several evidences supporting a coupling between morphological and chemical traits in roots (Fort, Jouany, & Cruz 2013; Prieto et al. 2015; Roumet et al. 2016; Marañón et al. 2020), some controversy at this respect still exists (Chen et al., 2013; Valverde-Barrantes, Smemo & Blackwood 2015; Weemstra et al. 2016; Kong et al. 2019). It is also worth noticing that we observed a strong relationship between SRA and root construction costs, but this relationship was weaker for SRL. This supports our previous evidence that SRA is more related to root tissue density than SRL in woody plants (de la Riva et al. 2018a), indicating that area-rather than length-based measurements could be a better indicator of the rate of investment per unit mass in roots. Our results evidence that the cost and energy involved in developing the structure of roots in woody species follow the trade-off between function and morphology determined by the RES (Roumet et al. 2016). Expanding this knowledge to a larger and more diverse pool of species worldwide would be necessary to determine whether these patterns may be generalized globally.

Our set of 60 woody species covered a wide range of fine root trait values within the root economics spectrum (RES); however, the observed range of root construction costs across species was rather low, supporting previous findings both for roots (Martínez et al. 2002; Villar et al. 2006) and leaves (Poorter & Villar 1997; Villar & Merino 2001; Villar et al. 2006). A possible explanation for these narrow ranges of CC is that trade-offs between chemical fractions and structural components tend to homogenize the construction costs across species (Poorter & Villar 1997; Martínez et al. 2002; Escudero, Mediavilla, Olmo, Villar & Merino 2017). For example, leaves in Mediterranean woody species with contrasting resource-use strategies differed slightly in their C concentration but allocated their C to different anatomical tissues; acquisitive species invested a greater proportion of C in metabolic tissues whereas conservative species invested a greater proportion of the C in structural tissues, which could be the case also for roots (de la Riva et al. 2016b; Escudero et al. 2017). This low variability in root construction costs could account also for the absence of significant differences across symbiotic association types in this study. In fact, both AM and EcM species showed different biomass stocks with similar carbon allocation rates in previous studies (Phillips & Fahey 2006; Valverde-Barrantes, Raich, & Russell 2007), which supports the lack of differences in construction costs in our set of species despite the observed differences in their morphological strategies (Fig. 1). Another possible explanation could be that roots carry out multiple functions, such as foraging ability, water transport, nutrient uptake and exchanging C and nutrients to support symbiotic associations, functions that may not always be necessarily maximized simultaneously (Comas et al. 2012; Bergmann et al. 2020). Whatever the case, the slight variation in construction costs across functional types in our study seem to respond to differences in the morphological traits of the RES.

We also showed that species segregation along the RES was conditioned by their type of symbiotic association (AM, EcM, ErM and N-fix). Species segregated along the RES gradient according to their main mycorrhizal associations; they varied from arbuscular mycorrhizal species (AM) having traits associated to more acquisitive strategies up to ectomycorrhizal species (EcM) in the other end, with traits associated to more conservative strategies. ErM species occupied an intermediate position and N-fixed species were distributed along all the whole gradient of resource economics. This segregation indicates that mycorrhizal types show affinity by a particular subset of morphological root traits (Kong et al. 2019), which allow them to persist under particular environmental conditions (Comas et al. 2012). Roots colonized by EcM fungi species normally present a higher lignification than those colonized by AM fungi (Brundrett 2002; Guo et al. 2008); a trait that is strongly associated to higher root tissue densities and root dry matter contents (Prieto et al. 2015; de la Riva et al. 2018a). On the other hand, roots from species colonized by AM fungi tend to exhibit trait values typical of acquisitive and fast-growing strategies, with a high investment in root area or length per unit root mass (i.e. SRA or SRL; Comas et al. 2012; Valverde-Barrantes, Smemo, Feinstein, Kershner & Blackwood 2018). One possible reason for these preferences could be that AM fungi species require specific anatomical and architectural adaptations to facilitate the root infection (e.g. lower root cortical areas and/or higher branching intensities; Comas et al. 2012). Such anatomical and architectural features increase the contact surface between the root and the AM symbiont (Valverde-Barrantes et al. 2018), and also enhance the root hydraulic conductivity and foraging ability of the host plant species, associated with a faster resource uptake, growth and development (Hernández, Vilagrosa, Pausas & Bellot 2009). Our results suggest that Mediterranean woody species that develop roots with acquisitive traits (e.g. high SRA) also favour the association with AM symbionts, resulting in a positive plant-symbiont feedback loop aimed at maximizing the plant C gain, which will benefit both the plant and the fungi (Allen & Allen 1991).

### Components of the root construction cost and symbiotic types

Despite similar relationships between root construction costs and the RES within groups of species with the same symbiotic association (Fig. 2), the influence of each component of the root construction costs on this relationship differed with the type of symbiotic association. These results suggest that the variation of the different components involved in root construction costs across species may be modulated by the specific resource uptake strategy of each symbiotic association type. The variation of root construction costs within roots of EcM species (situated on the conservative extreme of the axis) reflected a shift along their resource uptake strategy as a result of their different C and mineral concentration, which could suggest a different inversion on their structural components. Oppositely, the root from AM species (situated on the acquisitive side of the axis) seems to be mostly determined by a preferential investment on organic nitrogen, which is associated with higher concentrations of active metabolic compounds, such as proteins (Poorter & Villar 1997), while minerals and especially carbon concentration seems to exert a secondary role. This could be due to the fact that AM species are able to maintain their acquisitive performance with contrasting strategies: by investing carbon in cheap tissues that facilitate a faster return on investment, or by investing carbon in thicker roots that enhance the mycorrhiza association (trade-off from ‘do-it-yourself’ to ‘outsourcing’; Bergmann et al. 2020). In rhizobial N-fixing species, a similar proportion of the investment in construction costs was derived from C concentrations and organic nitrogen, possibly as a result of their wide distribution along the whole resource uptake gradient with both acquisitive and conservative species, which could lead to an interplay between structural and metabolic compounds according to economics spectrum expectations. We need to acknowledge here that this evidence is based on the species symbiotic preferences and, although tightly linked to root strategies, may not reflect direct measurements of root mycorrhizal colonization rates. Therefore, our conclusions on the different patterns of investment of root construction constituents in relation with symbiotic roles need to be validated in future studies by direct measurements of mycorrhizal fungi colonization. Nonetheless, we advocate that these findings represent a significant advance in the understanding of the relationship between root morphology and the resource optimization concept (Eissenstat & Yanai 1997).

### Variation of root construction costs along a soil resource gradient

Our prediction that variation of root construction costs across Mediterranean woody species would be related to the resource availability of their respective habitats was only partially supported. We found that root construction costs were influenced by resource availability and that the intrinsic cost (energy per unit of mass) of root production was higher in species growing in less fertile habitats than in those species inhabiting more favourable areas, supporting previous evidences using different woody species (Martínez et al. 2002) or different populations of *Quercus ilex* (Laureano et al. 2013). It has been frequently observed that soil productivity drives the functional trait composition on Mediterranean woody communities in line with the economics spectrum expectations (Cornwell & Ackerly 2009; de la Riva et al. 2018a). In this sense, the shrub species from the two communities in the Doñana National Park (the less fertile sites) showed the highest construction costs values (averages of 1.76 and 1.69 g glucose g^−1^ in Monte Negro and Monte Blanco, respectively). This was likely due to the dominance of conservative resource-use strategies in species inhabiting these sites, which has proven extremely advantageous to survive in these stressful environments (Lloret et al. 2016). This region is characterized by sand dune soils with very low water-retention, which explains its lowest fertility (Appendix S1). In line with these results, Martínez et al. (2002) found that the woody species from Doñana had higher root construction costs than others from more productive environments, due to their higher concentration of costly compounds, e.g. cell wall waxes that provide greater protection against water loss (Martínez et al. 2002; Samuels Kunst & Jetter 2008). By contrast, we found that in the most productive areas, such as riparian forests, dominant species were mostly deciduous with fast-growing strategies and low fine root construction costs (1.48 g glucose g^−1^, on average), probably in relation to less costly chemical compounds such as cellulose, and higher concentrations of minerals and organic acids, which allow them to achieve their highest growth rates (Poorter & de Jong 1999; Martínez et al. 2002). These results support the hypothesis that species from the most stressful environments need to expend comparatively more energy on root construction than do species under favourable conditions (Martínez et al. 2002; Villar et al. 2006; Laureano et al. 2013).

It is worth noticing that, in contrast with our expectations, we did not find a linear relationship between root construction costs and potential water availability, which may suggest an idiosyncratic pattern of carbon economics inversion along the gradient in water availability. This result is a priori surprising, not only because it is in contrast with the abovementioned findings in Mediterranean environments (Martínez et al. 2002; Laureano et al. 2013), but also because we have previously observed that root morphology was strongly associated with soil resource availability in Mediterranean woody species (de la Riva et al. 2018a). The main reason for this unexpected finding lies in the fact that the lowest construction costs (mean= 1.39 g glucose g^−1^) was found in plant species from the region of the gradient with the lowest water availability (Cabo de Gata-Nijar Natural Park). This site is extremely arid (de la Riva et al. 2018b) and, as shown in a previous study along a gradient of water availability in the Atacama Desert, root trait responses with aridity may not follow a linear pattern, and instead they present shifts at the most extreme ends (Carvajal, Loayza, Rios, Delpiano, & Squeo 2019). In the Atacama Desert, plant communities followed a complex pattern, whereby root traits shifted from acquisitive to conservative with increasing aridity, until a certain threshold in aridity was reached and species became more acquisitive again. Our results seem to follow this same pattern whereby fast resource uptake strategies are selected under very low water availability (i.e. Cabo de Gata), because they can survive by using the available nutrient resources during short periods of high water availability, e.g. after rain events (Chesson et al. 2004; Querejeta et al. 2018; Carvajal et al. 2019). Soils in Cabo de Gata were very shallow (de la Riva et al. 2018b) and plant species in this site have acquisitive root traits with very shallow root systems favoring fast, rather than slow, uptake strategies by faster water uptake immediately after rain events (Fort et al. 2013; Carvajal et al. 2019). That is, if the nutrient concentration is enough, these species are capable of faster and more profligate water and nutrient use to grow faster during short wet pulses. However, acquisitive plants usually have high nitrogen concentrations in their tissues to maintain high metabolic levels (de la Riva et al. 2016b), a nutrient that is frequently inaccessible in soils from arid ecosystems (Noy-Meir 1973; Ward 2009). Although specific mechanisms are challenging to infer, our results suggest that the role played by the type of plant symbiont association could be crucial to overcome this deficit. Associations with AM fungi and N-fixing bacteria often favour higher acquisition of nutrients (especially N) in low productive environments (Andrade et al. 2010; Smith & Smith 2011; Meng et al. 2015), thus supplying the necessary elements to maintain a higher metabolic activity and fast growth during these short periods of water availability. Indeed, the Cabo de Gata site had the highest proportions of AM species across all sites (with 61% of the species being AM) and one of the highest proportion of N-fixing species (22%), suggesting that symbiotic relationships may indeed be an advantage to maintain acquisitive strategies under resource scarcity, as previously hypothesized (Kramer-Walter & Laughlin 2018).

## CONCLUSIONS

This study supports that the differences in root construction costs in 60 woody Mediterranean species are a good reflection of differences in root morphological traits in line with expectations from the RES theory. We observed that different plant species have different strategies in their C investment that depend on their position along the resource gradient (RES) and on their main symbiotic association preference. The intrinsic components of the cost of root production also varied across species with contrasting symbiotic associations pointing to a trade-off between structural and metabolic compounds driven by the interplay between root economics and the type of symbionts. We also found that root construction costs are strongly modulated by soil resource availability (whether nutrients or water) following a non-linear pattern with water availability shifting from high to low construction costs at the most arid site, which points to a strong role of symbiotic associations in this shift. In summary, this study highlights that root construction cost is a fundamental parameter to understand the root resource strategies, although future efforts should be focused on clarifying the causes of different root CC values.

## ACKNOWLEDGEMENTS

This work was financially supported by the Spanish Ministry of Science and Innovation (Grants No. CGL2017-82254-R-INTARSU), ECO-MEDIT (CGL2014-53236-R), the project *Ecología funcional de los bosques andaluces y predicciones sobre sus cambios futuros* (For-Change) (UCO-27943) by Junta de Andalucía (Spain) and European FEDER funds and the Seneca Foundation (project 20654/JLI/18). We thank C. Navarro and C. Padilla for helping during field work, to the IRNAS Analytical Service for chemical analysis of plants and soil.

## AUTHORS’ CONTRIBUTIONS

E.G.R and R.V. conceived the ideas and designed the study; E.G.R, M.O., T. M and I.M.P.R conducted fieldwork; M.O. and R.V. conducted lab work; E.G.R. and I.P. performed statistical analyses; E.G.R and I.P. wrote the first draft and all the authors contributed significantly to revisions and gave final approval for publication.

