## Supporting material for "Root economics spectrum and construction costs in Mediterranean woody plants: the role of symbiotic associations and the environment"

**Supporting Information**

Appendix S1. Total number of woody species (Nº Sp) with the average of symbiotic association types (Arbuscular mycorrhizal species -AM-; Ecto mycorrhizal species -EcM-; Ericoid mycorrhizal species -ErM- and N-fixing species -N-Fix-), and mean data of the 7 environmental variables for site: organic matter (OM), soil nitrogen (SN) and availability of phosphorus (SP), magnesium (SMg), potassium (SK) and calcium (SCa), and potential soil water availability (PWA).

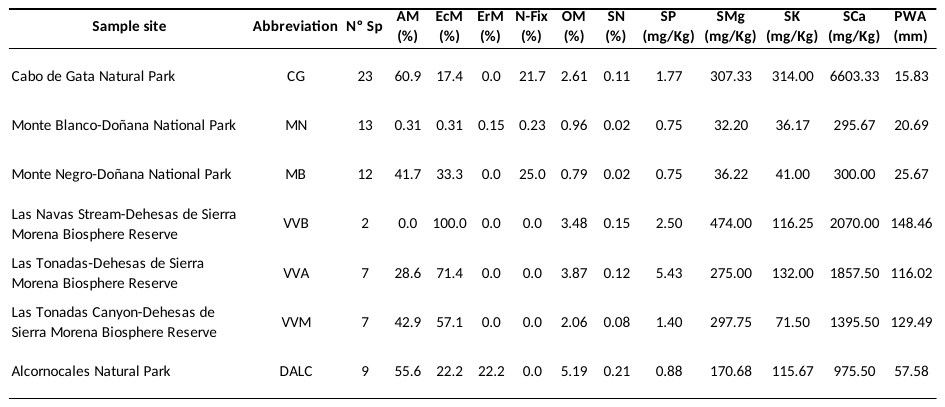

Appendix S2. List of the 60 woody species studied in south Spain. Species localization (sample site) and the main symbiotic association type are also indicated. See Appendix S1 for abbreviations of sample site.

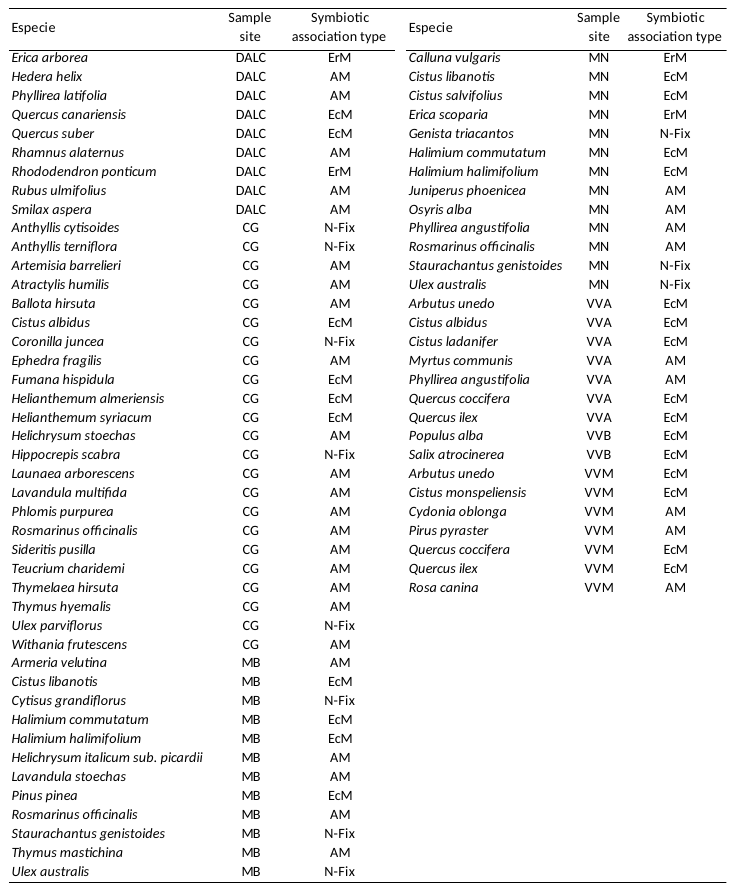

Appendix S3. Plot of the principal component analysis (PCA) with seven environmental variables (OM: organic Material; SP: soil phosphorus; SN: soil nitrogen; SMg: soil magnesium; SK: soil potassium; SCa: soil calcium; PWA: potential soil water availability)

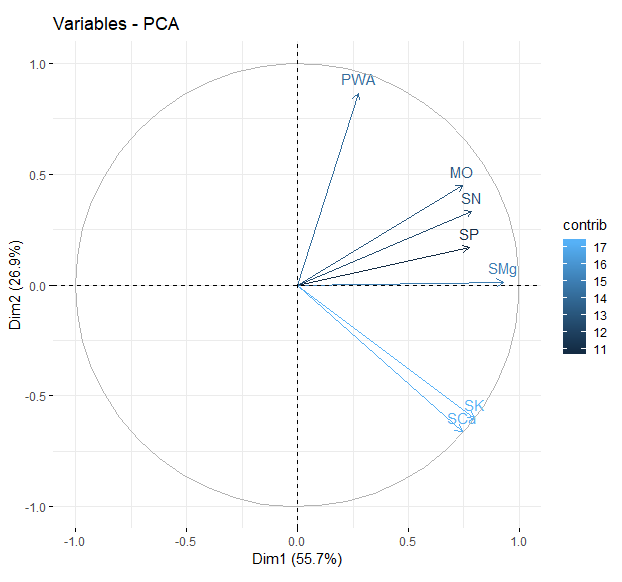

Appendix S4. The phylogenetic tree of the studied species, obtained with the PhytoPhylo tree (Qian & Jin 2016). Abbreviations of symbiotic association types: Arbuscular mycorrhizal species -AM-; Ecto mycorrhizal species -EcM-; Ericoid mycorrhizal species -ErM- and N-fixing species -N-Fix-.

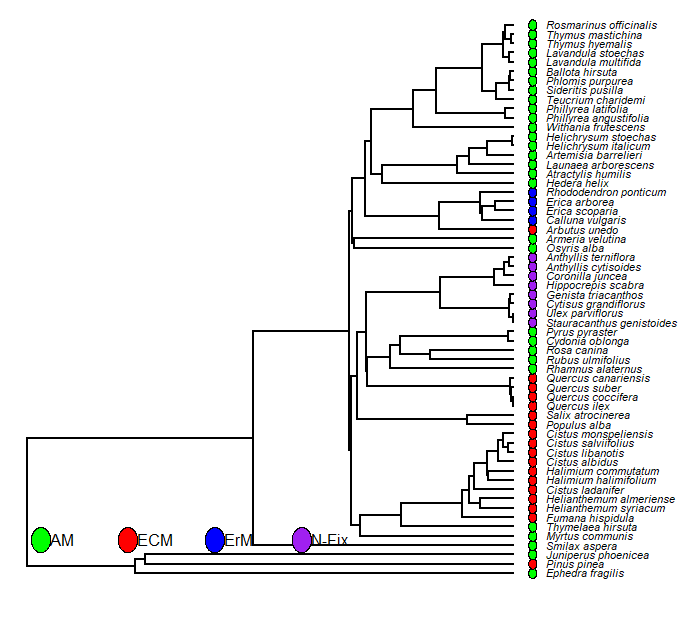

Appendix S5. Relationships between root construction cost and the second PCA axis (PC2) of root morphological traits depicting the trade-off between root density and diameter (R^2^=0.16; *P*=0.006).

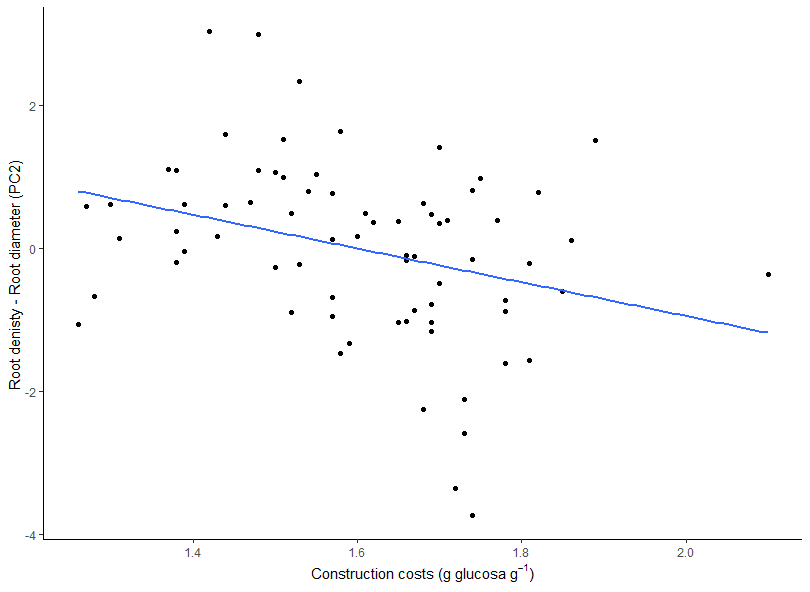
